# Within-infection diversity of *Plasmodium falciparum* antigens reflects host-mediated selection

**DOI:** 10.1101/212209

**Authors:** Angela M. Early, Marc Lievens, Bronwyn MacInnis, Christian F. Ockenhouse, Sarah K. Volkman, RTS,S Clinical Trial Partnership Committee of Investigators,, Dyann F. Wirth, Daniel E. Neafsey

**Affiliations:** Infectious Disease and Microbiome Program, Broad Institute of MIT and Harvard, Cambridge, MA, USA; Harvard T.H. Chan School of Public Health, Boston, MA, USA; GSK Vaccines, Rixensart, Belgium; PATH Malaria Vaccine Initiative, Washington, DC, USA; Simmons College, School of Nursing and Health Sciences, Boston, MA, USA

## Abstract

Host immunity exerts strong selection on pathogens, but it does not act in a uniform manner across individual hosts. By providing a direct approach for understanding host-specific selection pressures, patterns of intra-host pathogen diversity complement population genetic analyses performed on broad geographic scales. Here, we perform a combined analysis of inter- and intra-host diversity for the malaria parasite *Plasmodium falciparum* with haplotype sequences of three antigens sampled from over 4,500 natural infections in sub-Saharan Africa using targeted deep sequencing. We find that multi-strain infections in young children contain non-random combinations of parasite genotypes, and identify individual amino acid positions that may contribute to strain-specific blocking of infections. These results demonstrate for the first time that natural host defenses to *Plasmodium* detectably impact which infections proceed to the blood stage within a given host. This selection partially explains the extreme amino acid diversity observed at many parasite antigens and suggests that vaccines targeting such proteins should account for the impact of allele-specific immunity.

## Introduction

*Plasmodium falciparum*, the most deadly of the human malaria parasites, has been co-evolving with its human host for at least tens of thousands of years [1]. Over this time frame, the interactions of parasite, vector, and host have left signatures of strong selection on both human and *P. falciparum* genomes [2–4]. These evolutionary marks inform our understanding of how *P. falciparum* successfully invades multiple host and vector tissues while avoiding immune clearance. In turn, this knowledge can be used to develop novel ways of disrupting the *P. falciparum* transmission cycle. For instance, the high polymorphism of certain parasite proteins reflects their status as immune targets, suggesting that a vaccine composed of the same proteins could similarly elicit an effective immune response and impede infection.

Whole-genome sequencing on a population scale has used signatures of directional or balancing selection to identify immunogenic *P. falciparum* genes as potential vaccine candidates [5]. While these studies are useful for initially marking genes of interest, such population genetic analyses provide a relatively coarse-grained view of evolution; they identify the composite signals of diverse intra- and inter-host selection pressures, not selection directly mediated by individual vectors or hosts. To better inform therapeutic efforts, we would ideally decouple these various selection pressures by studying selective pressures imposed during different life cycle stages in isolation. In other pathogens, analyses of single infections have provided insights into within-host evolution [6,7], but such studies generally focus on *de novo* mutations arising during the course of a chronic infection.

Here, we describe a new high-resolution approach for studying the intra-host dynamics of *P. falciparum*, using high-depth sequencing of PCR amplicons and population genetic approaches that treat each infection as a discrete parasite population. In regions of high parasite prevalence, malaria infections often contain multiple haploid lineages of this sexually reproducing eukaryote. We investigate signatures of within-host selection on these parasite populations by using targeted deep population sequencing of three highly polymorphic genes that are known targets of B-cell or T-cell mediated immunity: *SERA2, CSP* and *TRAP* [8–11]. *SERA2* is a member of the serine repeat antigen gene family, and like its homologs, it is expressed during the disease-causing blood-stage of the parasite lifecycle. In contrast, *CSP* and *TRAP* are highly expressed by sporozoites [12], the form of the parasite that is injected into the human host by the mosquito and invades the host liver. Therefore, while all antigenic, these genes likely experience different selection regimes due to their divergent expression patterns across the *P. falciparum* life cycle (Fig 1).

**Figure 1.**
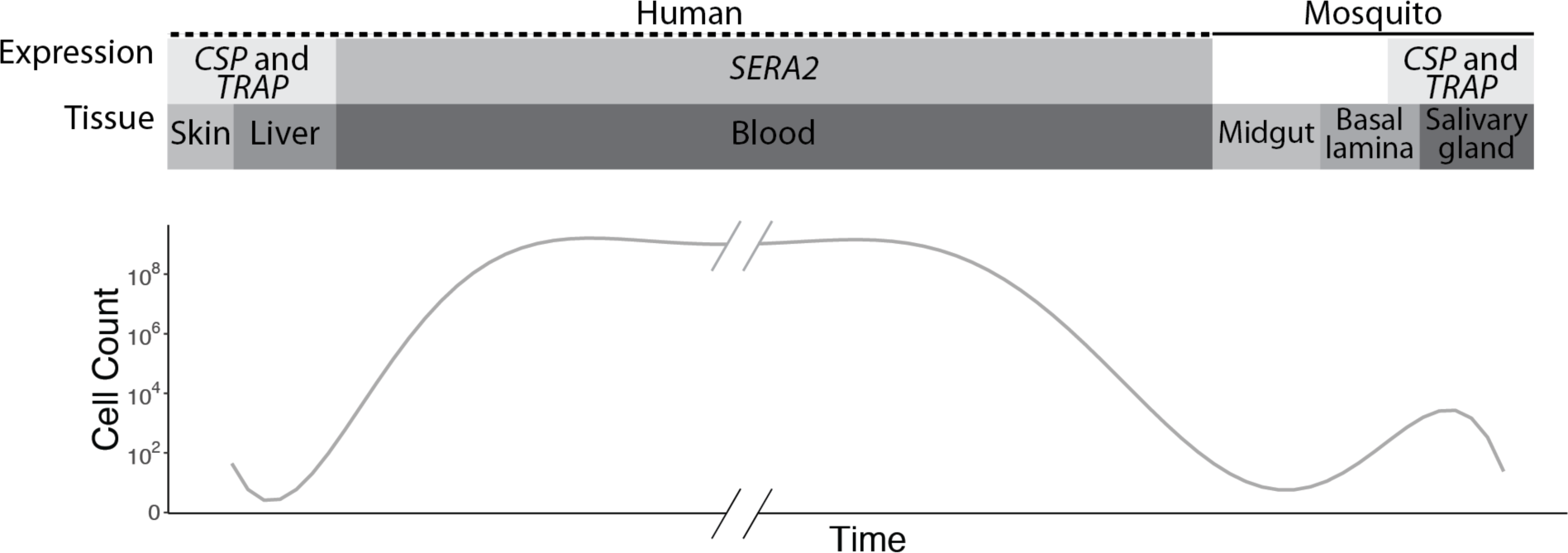
Schematic representation of *CSP*, *TRAP*, and *SERA2* expression profiles. The three genes have contrasting expression patterns that affect the source and duration of the selection pressures they experience.

Both CSP and TRAP are the focus of vaccine development, and several previous studies have examined their population-level diversity [13–22]. These loci exhibit high levels of nonsynonymous polymorphism in sub-Saharan Africa, and *TRAP,* in particular, has generally shown consistent evidence of strong selection (but see [19]). Several studies have documented functional evidence that natural and vaccine-mediated human immune responses differentially recognize variant CSP peptides [23–25]. Resolving the extent and nature of allele-specific immune recognition of these and other *P. falciparum* antigens would greatly aid the development of effective next-generation malaria vaccines. To date, however, observations of amino acid diversity have not been readily translated into an increased understanding of immune function. One complicating factor is that *CSP* and *TRAP* perform essential functions in both the mosquito vector and human host [26]. Ascribing selective pressures and functional constraint exclusively to the human host calls for an approach that decouples the selection imposed at these two distinct life cycle stages.

Immune selection on *P. falciparum* is complex; the parasite has developed numerous immune evasion strategies [27] and natural immune protection against malaria is mediated by recognition of a large number of antigens [28]. Dissecting host immune recognition with an evolutionary approach therefore requires a strategy capable of detecting relatively small selection coefficients through deep sampling of both the host and parasite populations. Multi-lineage *P. falciparum* infections are commonly characterized using small panels of putatively neutral SNPs [29], microsatellite markers [30], or size-length polymorphisms [31,32]. These approaches can quantify the number of clonal lineages within an infection [33,34], but are of limited use for understanding the mechanics of within-host selection. Whole genome shotgun sequencing has begun to provide information at the nucleotide level [35–37], but the predominance of host DNA in clinical samples limits the depth of parasite genome sampling. This means that low frequency variants within an infection are likely missed, constraining the use of these datasets to questions involving large selection coefficients.

In this study, we use deep sequencing of PCR amplicons to profile the parasite lineages within 4,775 infections, an approach that provides unprecedented power to detect selection events (blocked infections) with small effect sizes via the capacity to resolve the phasing of variants into haplotypes representing distinct lineages. The data show evidence of host-imposed selection occurring at both the sporozoite/liver stage and the blood stage of infection. Additionally, despite the large number of antigens that drive acquired immunity to malaria, we are able to point to specific amino acid positions that show a high probability of mediating allele-specific host recognition within two of these individual antigens, suggesting that the impact of acquired immunity on parasite fitness and infection outcome can be further dissected with high resolution genomic data.

## Results

### *CSP*, *TRAP*, and *SERA2* show patterns of diversity consistent with immune selection

Genes that experience immune selection are likely to exhibit high levels of nonsynonymous polymorphism and site frequency spectra that indicate the action of balancing selection. Both of these expectations hold for our three focal genes: *CSP*, *TRAP*, and *SERA2* (Table 1). To obtain a picture of these three genes in the context of genome-wide diversity patterns, we analyzed whole-genome *P. falciparum* sequence data from single-lineage infections in Senegal (n=99) and Malawi (n=110). All three of the focal genes are among the 5% most polymorphic genes in both populations, with *CSP* and *TRAP* being among the top 1%. The regions selected for amplicon sequencing display even higher levels of polymorphism than the full genes (Fig 2A). The site frequency spectra of these genes and amplicon regions are also skewed towards having a high proportion of intermediate-frequency SNPs, as measured by Tajima’s *D*, indicative of balancing selection. *TRAP*, as well as the regions of the *TRAP* and *CSP* genes chosen for amplicon sequencing, are within the upper 95^th^ percentile of both populations’ Tajima’s *D* distributions (Fig 2B). The genes, however, do not show evidence of extensive adaptation to specific geographic locations or host populations. None of the genes showed notable inter-population divergence as measured using *F_ST_*, and each amplicon-region showed lower population differentiation than its corresponding full gene (Fig 2C).

**Table 1.** Population genetic statistics for *CSP*, *TRAP*, and *SERA2* calculated using genome-wide sequencing data

|  |  | *CSP* |  | *TRAP* |  | *SERA2* |  | Genome-wide percentiles |  |  |  |  |
| --- | --- | --- | --- | --- | --- | --- | --- | --- | --- | --- | --- | --- |
|  |  | Full gene | Amplicon | Full gene | Amplicon | Full gene | Amplicon | 25 | 50 | 75 | 95 | 99 |
| Nucleotide Diversity | Senegal | 0.00601 | 0.0204 | 0.0102 | 0.0122 | 0.00191 | 0.00888 | 0.000218 | 0.000380 | 0.000634 | 0.00159 | 0.00399 |
|  | Malawi | 0.00806 | 0.0211 | 0.0102 | 0.0132 | 0.00198 | 0.00894 | 0.000225 | 0.000386 | 0.000625 | 0.00159 | 0.00396 |
| Nonsynonymous Nucleotide diversity | Senegal | 0.00674 | 0.0254 | 0.0120 | 0.0137 | 0.00181 | 0.0120 | 0.000144 | 0.000301 | 0.000534 | 0.00151 | 0.00385 |
|  | Malawi | 0.00713 | 0.0263 | 0.0120 | 0.0144 | 0.00197 | 0.0119 | 0.000154 | 0.000299 | 0.000528 | 0.00143 | 0.00425 |
| Tajima's *D* | Senegal | -1.39 | -0.132 | 0.536 | 0.298 | -1.21 | -0.494 | -2.21 | -1.91 | -1.52 | -0.381 | 0.548 |
|  | Malawi | -0.765 | 0.519 | 0.712 | 0.257 | -1.24 | -0.868 | -2.35 | -2.06 | -1.71 | -0.728 | 0.215 |
| *F<sub>ST</sub>* | Senegal-Malawi | 0.0512 | 0.0163 | 0.0576 | 0.0322 | 0.0236 | 0.00716 | 0.00769 | 0.0138 | 0.0255 | 0.0959 | 0.276 |

**Figure 2.**
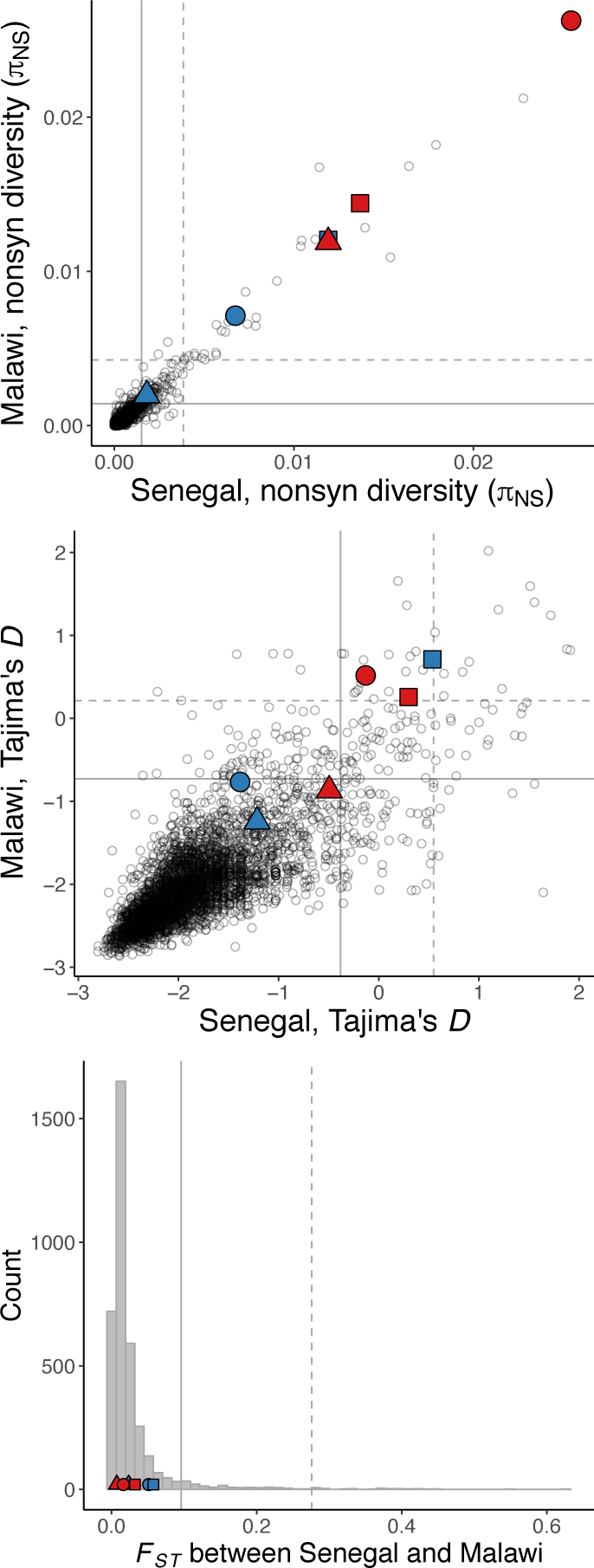
Genome-wide signatures of P. falciparum polymorphism in Senegal and Malawi. (A) *CSP, TRAP*, and *SERA2* show unusually high levels of nonsynonymous nucleotide diversity. *(B) TRAP,* the TRAP amplicon region, and the *CSP* amplicon region have unusually high values Tajima’s D. (C) None of the genes or amplicon regions show strong population differentiation (*F*_*ST*_). Each point represents the coding sequence from a single gene. Solid and dashed lines mark the 95^th^ and 99^th^ percentiles, respectively. Focal genes are marked with filled symbols: *CSP* (circle), *TRAP* (square), *SERA2* (triangle). Full genes are colored blue whereas the regions selected for amplicon sequencing are colored red.

PCR-based amplicon sequencing was conducted on 4,775 infections collected across five study sites in the context of a clinical vaccine trial (RTS,S [23]). Each amplicon was composed of approximately 300 nt of contiguous sequence, acquired from overlapping Illumina MiSeq reads. On average, thousands of reads were sequenced per amplicon per sample, providing a more fine-grained view of nucleotide diversity within the three focal genes. These data reveal a number of low frequency variants that were not detected in previous sequencing studies, but the overall patterns of polymorphism across the five study sites were consistent with what was observed in the Senegal and Malawi data (Table 2). The *CSP* amplicon contains three regions of high amino acid diversity [23] that include two previously described T-cell epitopes (Th2R and Th3R) [10] and a B-cell epitope (DV10) [9]. These epitopes physically co-localize to one side of the folded protein and surround a conserved hydrophobic pocket with extremely low amino acid diversity [38,39]. For the *TRAP* and *SERA2* amplicons there are no available protein structures, but their patterns of diversity are similar to *CSP*, showing discrete stretches of high diversity linked by regions with few or no variants (Fig 3).

**Table 2.** Summary of nucleotide variants described with targeted amplicon sequencing

| Amplicon region<br>(amplicon length) | Sequenced<br>amplicons | Total<br>variant sites | Nonsyn.<br>variant<br>sites | Singleton<br>variants | Common<br>variants* | Highly<br>variant<br>sites** |
| --- | --- | --- | --- | --- | --- | --- |
| ***CSP* (288 nt)** |  |  |  |  |  |  |
| Total | 3,821 | 55 | 50 | 15 | 23 | 12 |
| West Africa | 2,147 | 46 | 43 | 8 | 24 | 8 |
| East Africa | 1,674 | 41 | 39 | 8 | 23 | 8 |
| ***TRAP* (318 nt)** |  |  |  |  |  |  |
| Total | 8,774 | 52 | 52 | 13 | 15 | 5 |
| West Africa | 4,925 | 38 | 38 | 7 | 16 | 5 |
| East Africa | 3,849 | 38 | 38 | 10 | 15 | 3 |
| ***SERA2* (258 nt)** |  |  |  |  |  |  |
| Total | 9,007 | 67 | 67 | 19 | 12 | 16 |
| West Africa | 5,048 | 53 | 53 | 17 | 13 | 12 |
| East Africa | 3,959 | 49 | 49 | 11 | 8 | 11 |

\* minor allele frequency >0.02
\*\* >2 segregating alleles

**Figure 3.**
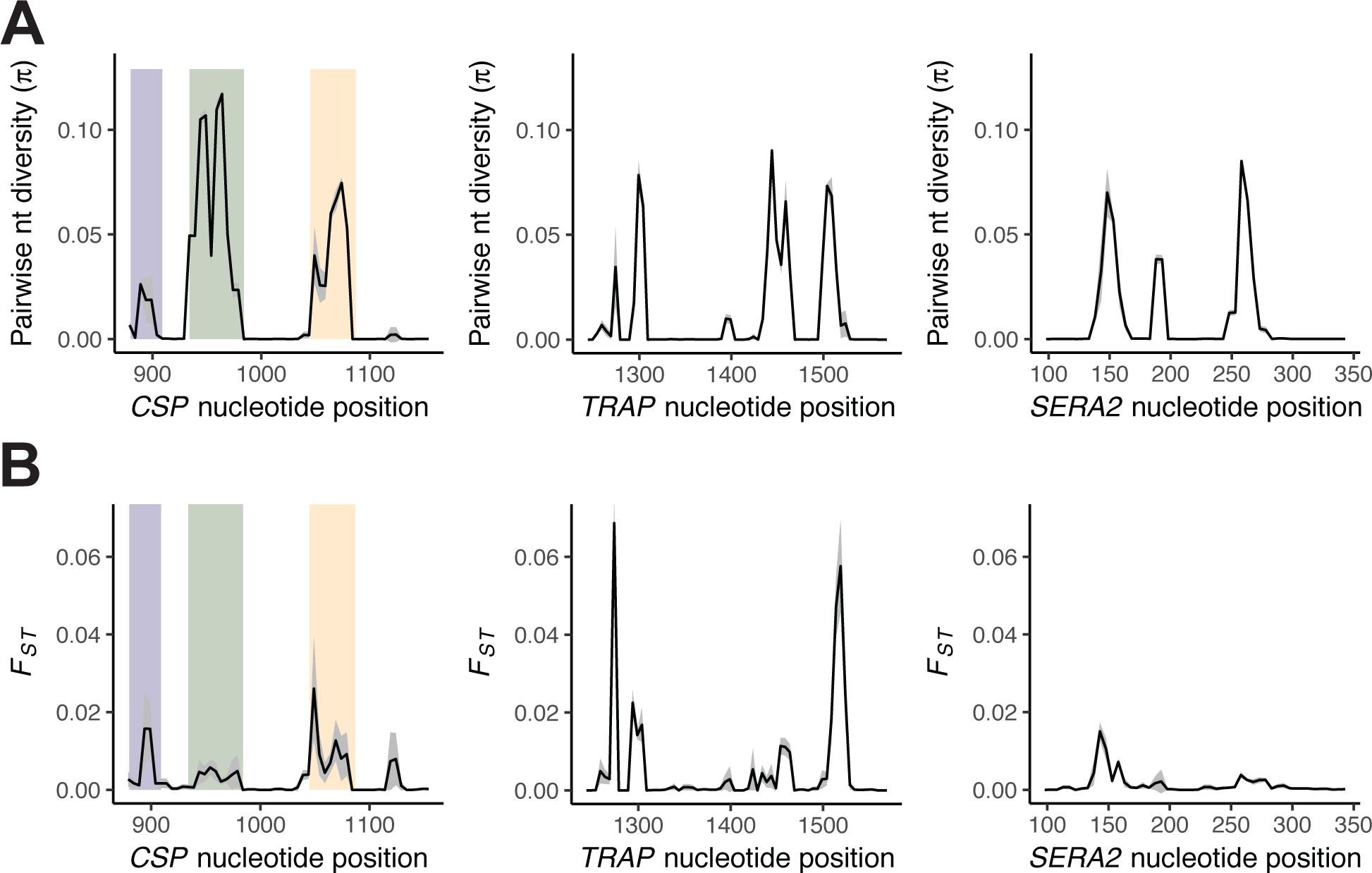
Nucleotide diversity and population differentiation within *CSP*, *TRAP*, and *SERA2* as measured with amplicon sequencing. (A) Graphs of pairwise nucleotide diversity (π) present the mean ± 1 S.D. across all five study sites. (B) Graphs of population differentiation show the mean ± 1 S.D. of pairwise *F*_*ST*_ values between western and eastern study sites. Statistics were calculated using SNPs within overlapping 10-nt windows with a step size of 5 nt. The colored bars in the *CSP* graph mark three previously identified regions of high diversity: DV10 (blue), Th2R (green), and Th3R (yellow).

As observed in the genome-wide data, population differentiation (*F*_*ST*_) between the western and eastern study sites is not unusually high for any of these amplicons, although there are a few individual variant sites that show reasonably high population differentiation (Supplementary Materials, Tables S1-S3). Interestingly, the two most diverse nucleotide sites in *CSP* (nt965 and nt952) show little population differentiation. These positions respectively contain four and three segregating nucleotides, and both have a pairwise nucleotide diversity (π) above 0.6 in all five study sites. Still, all population pairs show an *F*_*ST*_ under 0.01 at both positions suggesting they experience balancing selection at all geographic locations.

Because it provides haplotypic information, the amplicon sequencing approach also allows for an analysis of linkage disequilibrium (LD), or the preferential association of specific allelic variants on the same genetic background. We found that LD is highest for *CSP* and lowest for *SERA2* (Fig 4). Interestingly for *CSP*, LD exists both within and between known antigenic regions. While previously noted in the Th2R and Th3R regions, we found that this LD extends to the DV10 region as well. Previous work has found no evidence that intra-molecular forces drive this skewed association of alleles at specific amino acid positions [14]. This suggests that the maintenance of preferred allelic combinations within these positions might reflect the action of protein-protein interactions, which could include antibody recognition of conformational epitopes. Highlighting the unique nature of *CSP*, both *TRAP* and *SERA2* display lower levels of LD, suggesting that these proteins may experience fewer structural constraints or be subject to immune recognition through linear rather than conformational epitopes.

**Figure 4.**
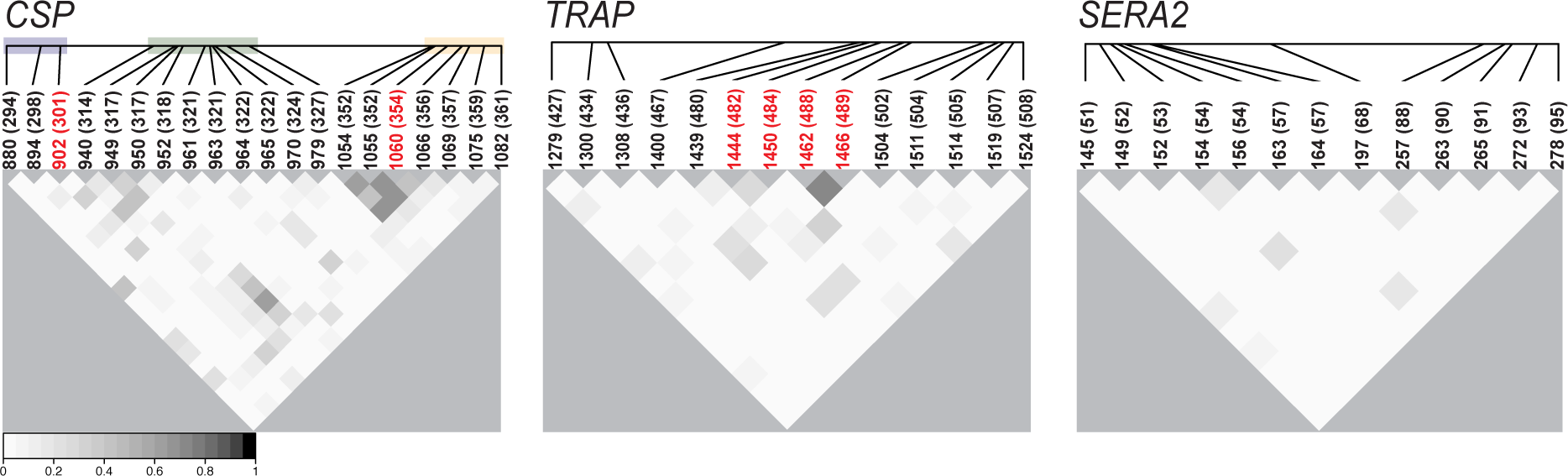
Linkage disequilibrium (LD) within the *CSP*, *SERA2*, and *TRAP* amplicon regions. LD between polymorphic nucleotides was calculated as Q*, an extension of r^2^ that allows for multiple alleles per site. Corresponding amino acid positions are in parentheses. Positions marked in red showed evidence for altered intra-host diversity at the amino acid level. For *CSP*, shading marks the positions within the DV10 (blue), Th2R (green), and Th3R (yellow) epitope regions. Only nucleotide positions with a major allele frequency <0.98 were included in the analysis. Data shown are from Nanoro, Burkina Faso. Observed LD trends were consistent across all five study sites (Supplementary Materials, Fig S1).

### Intra-infection diversity as a reflection of within-host selection

Patterns of LD—across *CSP* in particular—suggest that evolutionary forces are maintaining particular combinations of amino acids together on a single haplotype. Similarly, evolutionary forces may preferentially keep certain combinations of haplotypes together within a single infection. The majority of infections sequenced with the amplicon approach contained multiple haplotypes (*CSP*: 60.0%; *SERA2*: 53.6%; *TRAP*: 53.0%; Supplementary Materials Figure S2). This high rate of polygenomic infections allowed us not only to consider the population-level data described above, but also to conduct host-level analyses and look for evidence that amplicon haplotypes were distributed among infections in a non-random fashion. Such non-random associations would suggest that the host acts as a selective environment, lowering or enhancing a parasite’s fitness (*i.e.*, the likelihood of establishing a blood-stage infection) in a genotype-specific manner. This selection could result from selection within the observed infection, but also, as Hastings [40] demonstrated, from the cumulative effect of past selection events across multiple hosts as haplotype associations can be maintained through stable co-transmissions between individuals. We used simulations to model the expected distribution of haplotypes across infections and then compared the diversity in our observed infections to the diversity in these simulated infections (Fig 5). Non-random assemblages of observed haplotypes would cause a change in infection diversity in comparison to the neutral simulated scenario where infections occur in a completely random manner.

**Figure 5.**
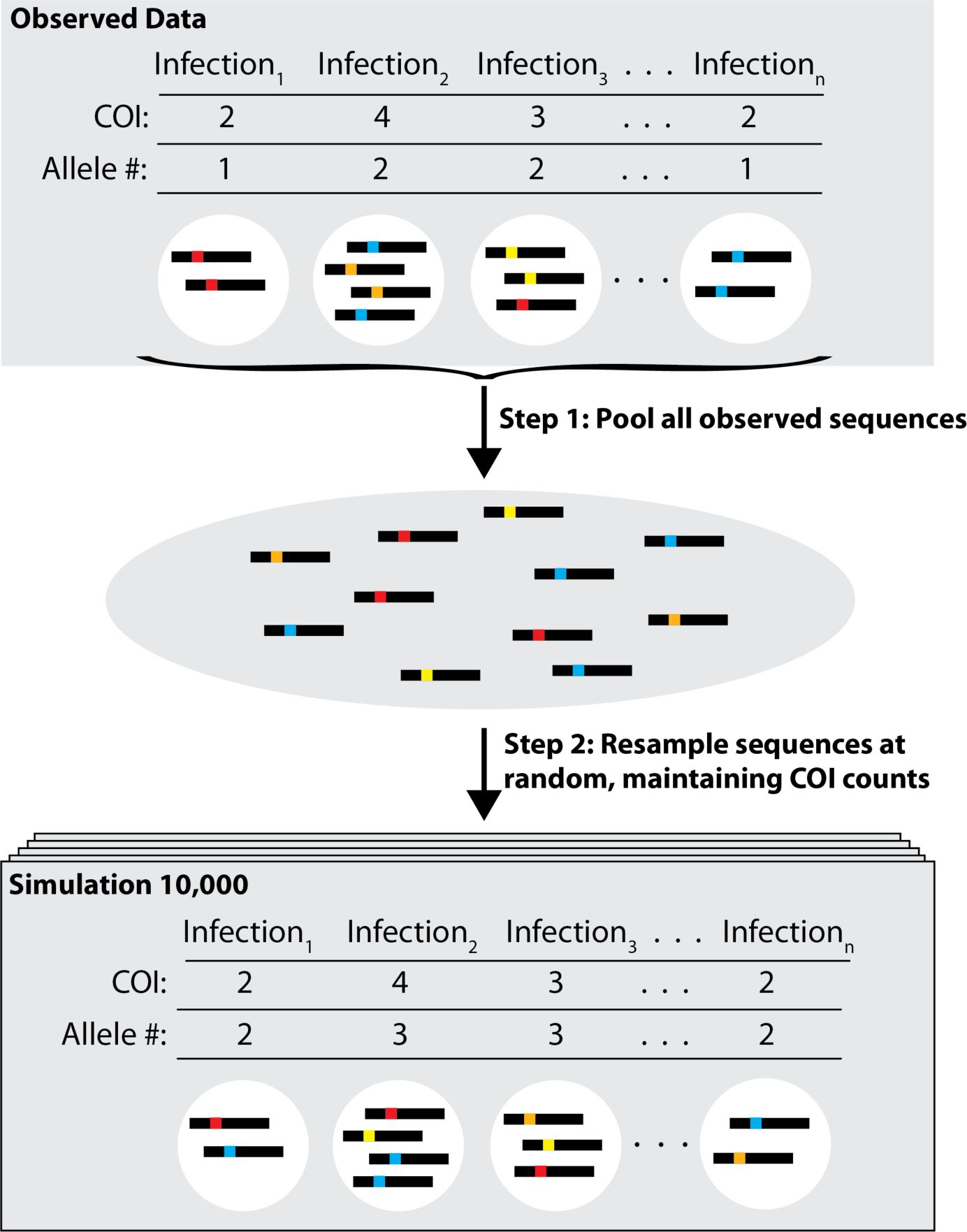
Infection simulations. For each sample location, 10,000 simulated datasets were created to test for the random distribution of haplotypes among infections. The number of haplotypes within each infection (COI) was held constant whereas the identity of each haplotype was varied through random sampling from the complete pool of sequenced amplicons. Within-infection diversity was then compared between the simulated and observed infections.

Observed within-infection amino acid diversity was lower than expected at all three genes across the five study sites (Student’s T-test with Fisher’s combined *P*, *P* = 0.0183 (*CSP*), *P* = 0.0123 (*TRAP*), *P* = 0.0219 (*SERA2*)). Selection, however, does not need to be evoked to explain this pattern; a reduction in observed heterozygosity is also expected under a model of population sub-division (the Wahlund effect). While we earlier observed that population differentiation is relatively low between study sites (Fig 3), *P. falciparum* populations are known to contain microstructure at the level of a household or village [41], which could contribute to protein-wide reductions in within-infection diversity. However, while population sub-structure would cause an overall decrease in heterozygosity across the given protein, the change in heterozygosity at any individual site would be random. We therefore looked for amino acid variants that showed decreased within-infection heterozygosity across all five geographic sites. This pattern would suggest the consistent action of selection rather than the random effects of genetic drift operating on structured sub-populations.

Across all three amplicons, five amino acid positions showed statistically significant reductions in diversity (Fig 6). Interestingly, the amino acid positions identified in CSP (aa354) is one of the seven amino acid positions that showed significant differential protective efficacy in a phase 3 trial of the malaria vaccine RTS,S [23]. In Neafsey *et al*. [23], this was observed as a vaccine-induced effect that manifested as greater protection against infection by parasites harboring a CSP haplotype matching the vaccine construct in RTS,S-vaccinated individuals. In this study, we limit our analysis of CSP to individuals who were enrolled in the control arm of the trial and therefore only detect the effects of natural immunity. Combined, these studies provide two independent lines of evidence suggesting that this residue is recognized in an allele-specific manner by an adaptive B- and/or T-cell response.

**Figure 6.**
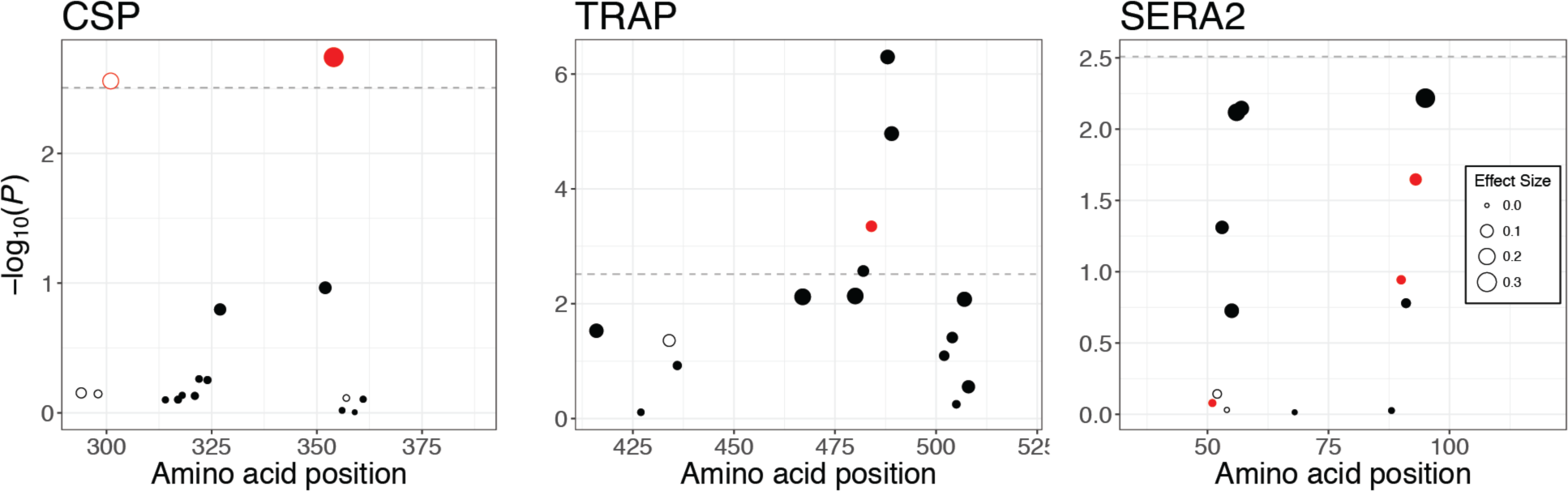
Deviation from expected within-host diversity at individual amino acid positions. Points above the dotted line mark amino acid positions with significant heterozygosity differences between observed and simulated infections after Bonferroni correction. Filled circles show a reduction in observed within-infection diversity whereas open circles show an increase in observed within-infection diversity. The size of the point corresponds to the effect size estimated with a quasi-Poisson regression. Points in red show a marginally significant effect of age on diversity (*P*< 0.05) before multiple testing correction, but none of these interactions remain significant after Bonferroni correction. Only variant positions with a major allele frequency <0.98 were included in the analysis.

Given past evidence that specific haplotypic combinations enhance parasite infectivity [42], we also considered the inverse question: do certain genotypes preferentially associate within infections? Only one amino acid position (CSP aa301) showed evidence of increased within-infection heterozygosity. As with CSP aa354—which showed increased within-host homozygosity—CSP aa301 was identified by Neafsey *et al.* [23] as showing an effect of allele-specific vaccine efficacy in the RTS,S vaccine trial. One of the nucleotide positions coding for this amino acid (nt902) is also the *CSP* nucleotide position with the highest divergence (*F_ST_*) between western and eastern study sites. Overall, however, the relative low number of sites showing increased heterozygosity suggests that immune interference does not widely contribute to changes in infection diversity—at least not at the level of individual amino acid variations. Instead, the stronger evolutionary force in these populations appears to be allele-specific selection against particular genotypes, leading to an overall decrease in within-infection diversity.

### Patient age negatively correlates with intra-host diversity at a blood-stage antigen

The above analyses suggest that either the host or mosquito vector serves as a selective barrier, causing infections to be non-randomly assembled. To further test the hypothesis that selection is specifically exerted by adaptive immune responses, we investigated how infection diversity relates to a known correlate of immune competence: patient age. In malaria-endemic countries, adults do not acquire full immunity to malaria infection [43], but the parasite levels in their blood are significantly reduced compared to the levels in young children, and they are less likely to develop clinical malaria [44]. The acquisition of adaptive immune defenses plays a role in this age-dependent disease resistance [45], although the exact mechanism remains unclear [39]. Our dataset offers an opportunity to examine the effects of age-related immune acquisition on a nucleotide level. To do so, we make two assumptions: (1) hosts mount more effective immune responses against parasite haplotypes to which they were previously exposed; and (2) patient age positively correlates with cumulative exposure to *P. falciparum*. Under this model, successful infection blocking would be haplotype specific and more frequent in older relative to younger children. This would lead to lower infection diversity in the former relative to the latter group at loci that are effective immune targets.

To test if patient age correlates with *P. falciparum* infection diversity, we constructed a quasi-Poisson model that accounted for additional factors including study site, complexity of infection, and type of sampling (active versus passive). For SERA2, but not CSP or TRAP, we found a negative effect of patient age on within-infection amino acid diversity (-0.032 mismatches per year, *P* = 0.028; Table 3). Across the SERA2 amplicon, mean number of pairwise differences per infection was 2.907. This suggests that as children age by one year, the amino acid diversity at SERA2 reduces by approximately 1.1%.

**Table 3.** Parameter estimates from quasi-Poisson regression model of within-host infection diversity

| Variable | CSP |  | TRAP |  | SERA2 |  |
| --- | --- | --- | --- | --- | --- | --- |
|  | Effect Estimate | *P* | Effect Estimate | *P* | Effect Estimate | *P* |
| Patient age (in years) | 0.0094 | n.s. | 0.015 | n.s. | -0.032 | 0.028 |
| Sample Type* | -0.033 | n.s. | 0.016 | n.s. | 0.072 | 0.0015 |
| Complexity of Infection** | 0.011 | n.s. | -0.012 | 0.035 | -0.0020 | n.s. |

\* longitudinal, asymptomatic sampling vs clinical infection sample
\*\* number of distinct observed haplotypes within the infection

We next modeled diversity at individual amino acid positions within each amplicon. Diversity at two amino acid positions in the *SERA2* amplicon (aa93 and aa95) showed nominally significant age effects (aa93: -0.12 mismatches per year, *P* = 0.047; aa95: -0.39 mismatches per year, *P* = 0.032), but their statistical significance did not survive multiple-testing correction. Overall, therefore, this result suggests that the reduction in amplicon-level diversity cannot be attributed to the effect of a single amino acid site. This is consistent with a model where immune recognition at this antigen occurs across multiple sites rather than at a single residue.

Given that *SERA2* is expressed during the protracted blood-stage, while *CSP* and *TRAP* are both expressed during the shorter sporozoite/liver-stage, this observation is consistent with evidence that natural acquired immunity most strongly controls *P. falciparum* transmission at the blood-stage of infection [47], although our sample size of three genes is too small to draw robust conclusions from this pattern. Detection of this trend in young children (< 5 years of age) suggests that acquired immune defenses likely begin building at an earlier age than previously observed in clinical studies [47]. These defenses, however, may not yet be sufficient to cause reductions in infection that manifest in patterns of reduced disease incidence in older children.

## Discussion

Targeted deep sequencing of pathogen genes provides a powerful and efficient way to gain insight into instances where genes experience diverse or conflicting selection pressures. Our results with *CSP*, *TRAP*, and *SERA2* demonstrate that population-level and host-level approaches provide complementary information that together identify not only genes but also specific amino acid positions of evolutionary and immunological interest. Gene-level statistics can mask relevant intra-gene patterns, especially when genes include constrained functional elements. Similarly, population-level analyses average selection pressures across multiple life stages. By focusing on a single point in the life cycle, we find evidence for there being amino acid positions under selection within *CSP* and *SERA2—*genes that exhibit non-significant evidence for selection in a traditional population-level genome-wide analysis. In this regard, there are parallels between the analyses we present here and approaches such as selection components analysis [48] and single-cell sequencing. In essence, these are all ways of dissecting a single composite signal to better understand the distinct forces or dynamics underneath. This added granularity helps assign causality and identify loci targeted by selection that may be masked by averaging signals across loci or life stages.

The amino acid positions we identify with this approach may mediate allele-specific immune responses to *P. falciparum*. While we do not know the extent to which parasite antigens are or are not cross reactive, several recent vaccine candidates—all monovalent subunit vaccines developed from single *P. falciparum* genotypes—have displayed allele-specific protection [23,49]. In light of these observations, one proposed course of action is the development of multivalent vaccines that target multiple pathogen genotypes [50]. By identifying specific amino acid positions that likely contribute to differential recognition, we provide key information for the development of such next-generation vaccines. Further, we illustrate that the challenges facing vaccine development with the leading vaccine component (CSP) differ from those confronting vaccines that would use antigens like TRAP and SERA2. Due to a combination of protein function and expression pattern (Fig 1), each protein experiences distinct selection pressures and structural constraints that create unique signatures of diversity, divergence, and LD.

While previous genetic analyses have provided evidence that *CSP* is under selection in the human host [13–17], few have provided actionable information on specific amino acid positions that may mediate natural host-pathogen interactions. One important exception is Gilbert *et al.* [42], who studied within-host haplotype diversity of the CSP T-cell epitope Th3R. Like this study and Neafsey *et al.*[23], they found that alleles at CSP amino acid 354 show preferential associations within infections. Interestingly, however, their samples showed the opposite effect of the one observed here. In our study, diversity at aa354 is lower within hosts than would be expected. Conversely, Gilbert *et al.* [42] found that two 8-aa haplotypes (cp26 and cp29) that differed only at aa354 showed the propensity to co-occur within infections, thereby increasing diversity at this single amino acid position. Additional *in vitro* studies found that these two haplotypes acted in concert to evade immune detection through a process of immune interference, thereby increasing their combined fitness [51]. In our samples, one of these haplotypes was at too low of a frequency to assess its potential preferential associations. Thus, the diversity changes we observed at aa354 were independent of the 8-aa haplotypes described by Gilbert *et al.* [42]. The opposing directions of the effects noted across studies suggests that non-additive interactions between alleles can determine the course of selection on any individual site, reinforcing the need to consider extended haplotype information and patterns of LD when studying immune recognition of *P. falciparum* antigens.

As we demonstrate here, targeted deep sequencing is a powerful and efficient method for gaining insight into immune interactions. The utility of this approach, however, extends beyond questions of immune selection. There is growing recognition that intra-host dynamics must be considered when studying diverse aspects of *Plasmodium* biology, including drug resistance [52,53], disease severity [54–57], and transmission rate [58,59]. Our observations show that human-mediated selection on *P. falciparum* is sufficiently strong to be detected using measures of within-infection diversity and demonstrate the utility of incorporating evolutionary approaches into *Plasmodium* studies. Targeted deep sequencing of genes involved in additional biological processes can further augment current efforts to understand and manipulate the interactions between humans and this complex parasite.

## Methods

### Genome-wide population genetic data: filtering and analysis

To place our amplicon sequences within a genome-wide context, we obtained genome-wide variant calls for samples collected in Senegal [60] and Malawi [20] from the Pf3k project (release 5; www.malariagen.net/projects/pf3k). Variant calls were downloaded as vcf files, and only variable sites that were flagged as passing the GATK VQSLOD filter were retained. We further limited the analysis to variant sites that fell within coding regions and used SnpEff calls to annotate each variant as synonymous or non-synonymous. For 5,077 genes, we calculated the percentage of variant calls that were heterozygous or missing for each sample. We identified a set of 243 genes where 20% or more samples had either >=25% heterozygous calls or >= 25% missing calls. After masking these genes, we calculated the fraction of variant sites across all genes that were heterozygous or missing for each sample. *Plasmodium* is haploid throughout its human stages. Heterozygous calls therefore signal the presence of multiple co-occurring genetic lineages within a sample. Samples with either >= 2% heterozygous calls or >= 4% missing calls were removed from the analysis. This left us with a set of 99 Senegal and 110 Malawi samples that likely contained a single *P. falciparum* genotype (Supplementary Materials, Table S1). We further removed 315 genes that had either >=20% heterozygous calls or >= 20% missing calls in these remaining samples. We transformed any remaining heterozygous calls into homozygous calls by retaining only the allele with greatest read support. Only SNP calls were retained for use in downstream analyses.

For each gene, we calculated two measures of nucleotide diversity (π and Watterson’s θ), Tajima’s *D*, and Weir and Cockerham’s estimate of *F*_*ST*_ [61] using custom Perl scripts. Sites with more than two alleles were retained and counted as multiple segregating sites when calculating Watterson’s θ. The numerator and denominator of the *F*_*ST*_ statistic were averaged individually across sites in a given gene. At each site, we adjusted the sample size based on the number of samples with successful calls. We assumed that all sites without called variants were reference alleles for all the samples. For each gene, statistics were calculated across all sites, at synonymous sites only, and at non-synonymous sites only. To ensure we were not biasing our results by using genes with little coverage, we limited our analysis to 3,682 genes that had at least five SNPs in one of the populations.

### Sample collection, sequence generation, and error filtration

The amplicon sequences for *CSP* (PF3D7_0304600), *TRAP* (PF3D7_1335900), and *SERA2* (PF3D7_0207900) were previously generated by Neafsey, *et al.* [23]. The *CSP* amplicon covers the diverse C-terminal region of the *CSP* gene, which forms a large portion of the RTS,S vaccine construct (Pf3D7_03_v3:221352..221639) [62]. The *TRAP* and *SERA2* amplicons were designed to capture regions of high diversity within these respective genes (Pf3D7_13_v3:1465059..1465376; Pf3D7_02_v3:320763..321020). Parasite DNA was obtained from blood samples acquired as part of the RTS,S vaccine phase 3 trial [63]. In brief, children (5 to 7 months of age) and infants (6 to 12 weeks age) were randomly assigned to one of two arms of the vaccine trial. Approximately two-thirds received the RTS,S/AS01 vaccine regimen whereas one-third received a control rabies vaccine. Blood samples included both passive and active sampling. Passive clinical samples were collected from febrile children with at least 5000 parasites/mcl at the time of first clinical malaria infection occurring within the first 12 months following completion of the triple-dose vaccination regimen. Active cross-sectional sampling occurred in a subset of participants at 18 months post-vaccination. Samples were stored as dried blood spots until DNA extraction. Targeted regions were then amplified and sequenced using overlapping 250-bp paired-end reads with Illumina MiSeq technology. Paired-end reads were merged using FLASH [64] and aligned to the PlasmoDB v.9.0 3D7 *P. falciparum* reference genome using BWA v.0.74 [65].

This targeted amplicon approach allowed the identification of complete, linked haplotypes across the sequenced regions. To minimize the misidentification of spurious haplotypes within samples, Neafsey, *et al.* [23] adopted a set of conservative filters that removed the following: (1) all reads with uncalled bases; (2) sequences representing fewer than 1% of the reads in a given sample; (3) haplotypes observed only once in the entire dataset that were represented by fewer than 500 reads; (4) sequences that showed evidence of PCR chimerism. Further, haplotypes within each sample were clustered. If two haplotypes differed by only 1 nucleotide (nt) and their read ratio was less than 0.15, the haplotypes were merged. After merging and filtering, the median read count per sample was still very high (*CSP*: 5559, *TRAP*: 3001, *SERA2*: 4912).

For the purposes of this study, we used data from the five largest study locations: Kintampo, Ghana; Agogo, Ghana; Nanoro, Burkina Faso; Kombewa, Kenya; and Siaya, Kenya. In total, this provided information on 4,775 infections: 2,854 from RTS,S-vaccinated and 1,921 from control-vaccinated individuals. Analyses of *CSP* used only data from the control-vaccinated individuals. After confirming that the RTS,S vaccine had no effect on diversity at the *TRAP* and *SERA2* loci, we included all patient samples in analyses of these loci.

### Population genetic and statistical analyses

For each study site, we calculated pairwise nucleotide diversity (π) and Weir and Cockerham’s measure of population differentiation (*F_ST_*; [61]) at each SNP and within 10 nt windows across each gene amplicon. We calculated linkage disequilibrium (LD) between pairs of sites—including sites with more than two variants—using Lewontin’s D’ [66] and Hedrick and Thomson’s Q* [67].

To test for the effect of age on within-infection amino acid diversity, we constructed a quasi-Poisson regression model with patient age at time of infection, sample site, sample type (passive/clinical, active/cross-sectional), and complexity of infection (COI; the number of genetically distinct clonal lineages within one infection) as fixed effects. Within-infection diversity was modeled separately for each amplicon. Models were constructed to study infection diversity on the level of the whole amplicon and on the level of individual amino acids. The analysis was conducted in R with the glm function.

### Modeling expected infection heterozygosity

Within our sample of 4,775 infections, we recovered 0-13 haplotypes per amplicon per infection, with over 50% of infections containing multiple segregating *P. falciparum* genotypes. Infection heterozygosity is dependent on COI, and so expected heterozygosity cannot be calculated as for a diploid organism. Instead, we determined the expected level of infection heterozygosity through bootstrapping. For each sample site, we created a haplotype pool that contained the exact number of each haplotype recovered during sequencing. We then created 10,000 sets of simulated infections that maintained the same COI structure observed in each population but where the infecting strains were drawn randomly (with replacement) from the haplotype pool (Fig 5). As our sequencing is not able to differentiate identical haplotypes, each simulated infection was constrained to contain only unique haplotypes.

For each sample (both observed and simulated), we calculated within-infection amino acid diversity for the full length of each amplicon as the mean number of differences between each pairwise haplotype comparison within the sample. When analyzing diversity at individual positions, we counted the number of haplotype pairs that mismatched at the given site. To determine whether mismatches were more or less frequent in the observed vs. the simulated infections at a given amino acid position, we constructed a quasi-Poisson regression model that included data set (observed, simulated), COI, study site, sample type (passive/clinical, active/cross-sectional), and age group (infant, child) as fixed effects. We additionally included two interaction terms: (data set x sample type) and (data set x age group). These accounted for the possibility of increased acquired immunity in older children and for potential differences between symptomatic and asymptomatic infections. The Bonferroni correction was performed to account for multiple testing across multiple sites within each individual gene. Variant sites were only analyzed if they were polymorphic in all five populations and had a global major allele frequency under 0.98. For the simulated data, we included results from 1,000 bootstrapped data sets. The analysis was conducted in R with the glm function.

## Data availability

All raw Illumina reads from the original Neafsey *et al.* [23] study are available on the NCBI Short Read Archive (BioProject PRJNA235895). Supplementary Tables S5 and S6 contain the final called haplotypes and metadata from children and infants enrolled in the control arm of the vaccine trial.

## Authors’ contributions

AME and DEN conceived of the analysis. AME conducted the analysis and drafted the manuscript. DEN and DFW provided funding and project guidance. BM and SKV provided project administration and oversight. PBG, ML, CFO, SKV, DFW, and DEN contributed to the development of original clinical study. All authors reviewed the manuscript and gave final approval for publication.

## Competing interests

ML is an employee of GSK group of companies, which is involved in malaria vaccine development. All other authors declare no competing interests.

### Acknowledgements

We thank the participants of the RTS,S phase 3 clinical trial and their parents as well as the members of the RTS,S Clinical Trials Partnership. Aimee Taylor, Seth Redmond, and members of the Neafsey and Wirth research groups provided thoughtful comments and discussions.

## Supplementary Material

**Table S1: Nucleotide diversity and population differentiation for variants in the CSP amplicon**

**Table S2: Nucleotide diversity and population differentiation for variants in the TRAP amplicon**

**Table S3: Nucleotide diversity and population differentiation for variants in the SERA2 amplicon**

**Table S4: List of Pf3k samples used in analysis**

**Table S5: Unique CSP, TRAP, and SERA2 haplotypes found in samples collected from children enrolled in the control arm of the RTS,S/AS01 phase 3 vaccine trial**

**Table S6: Haplotype IDs and select associated metadata for samples collected from children enrolled in the control arm of the RTS,S/AS01 phase 3 vaccine trial**

**Figure S1. LD plots for nucleotides within the CSP, TRAP, and SERA2 amplicon regions for each of the five study sites**

**Figure S2. Number of haplotypes sequenced per infection for each amplicon region.**

